# The cowboy effect: robot gaze influences human decisions

**DOI:** 10.1101/2020.10.21.345876

**Authors:** Marwen Belkaid, Kyveli Kompatsiari, Davide de Tommaso, Ingrid Zablith, Agnieszka Wykowska

**Author notes:** Equal contribution (Marwen Belkaid, Kyveli Kompatsiari). **Corresponding author information:** Agnieszka Wykowska, Istituto Italiano di Tecnologia, Center for Human Technologies, Via Enrico Melen, 83, 16152 Genoa, Italy, (AW). **Email addresses**, (MB), (KK), (DDT), (I.Z.).

## Abstract

In most everyday life situations, the brain needs to engage not only in making decisions, but also in anticipating and predicting the behavior of others. In such contexts, gaze can be highly informative about others’ intentions, goals and upcoming decisions. Here, we investigated whether a humanoid robot’s gaze (mutual or averted) influences the way people strategically reason in a social decisionmaking context. Specifically, participants played a strategic game with the robot iCub while we measured their behavior and neural (EEG) activity. Participants were slower to respond when iCub established mutual gaze prior to their decision, relative to averted gaze. This was associated with a higher decision threshold in the drift diffusion model and accompanied by more synchronized EEG alpha activity. In addition, we found that participants reasoned about the robot’s actions in both conditions. However, those who mostly experienced the averted gaze were more likely to adopt a self-oriented strategy and their neural activity showed higher sensitivity to outcome. Altogether, these findings suggest that robot gaze acts as a strong social signal for humans, modulating response times, decision threshold, neural synchronization, as well as choice strategies and sensitivity outcomes. This has strong implications for all contexts involving human-robot interaction, from robotics to clinical applications.

## Introduction

Many human decisions are made in social contexts, which often requires assessing the intentions of others. In order to infer others’ mental states and to be able to predict their behavior, humans rely largely on nonverbal cues. In particular, eye contact is a strong communicative signal in human interactions which can convey information about one’s state, goals, intentions, or willingness to interact (Kleinke, 1986; Emery, 2000). The effect of mutual gaze has been extensively investigated in studies showing that gaze affected arousal level (for a review see Hietanen et al., 2018) as well as various cognitive processes (including memory, attention, and motor actions; for a review see Hamilton (2016)). However, the specific effect of gaze in the context of social decision-making has been rarely addressed, and certainly not in the context of human-robot interaction.

Social interactions rely to a great extent on efficient cooperation with other humans. However, every decision to cooperate with others entails the risk of being exploited. Behavioral and neuro-economics have been studying complex social dynamics using tasks adapted for game theory. These strategic games have been specifically designed to address social decisions in conflicting situations which may involve coordination, reciprocity, risk-taking, competition, altruism, and other mechanisms. In particular, the “Chicken game” (Rapoport & Chammah, 1966) depicts a situation in which two drivers move towards each other on a collision course: one must yield (deviate), otherwise both cars crash. If only one of them yields, this player is called a “chicken” (i.e. coward). In this game, cooperation (i.e. when both deviate) leads to the highest joint payoff while unilateral cooperation (i.e. when only one deviates) is disadvantageous. Competitive approach (driving straight towards the other player) generates the maximal individual payoff, if it is unilateral, but with the risk of a high punishment in case the other player decides for the same action.

The chicken game has been previously employed in human neuroscience research; for instance, to investigate the influence of personal traits (Wang et al., 2017) or dyad familiarity (Chen et al., 2017) on participants’ behavior and neural activity. More generally, various studies have combined strategic games with neuroimaging techniques in the quest for a better understanding of the neural mechanisms at play in social decision-making (see Rilling & Sanfey (2011) for a review). However, these studies are generally screen-based and lack the ecological validity of real-time interactions. In this regard, robots offer a powerful solution for the design of controlled, yet naturalistic and embodied, interactions. Previous works in human-robot interaction have also proposed experiments based on strategic games, comparing different types of robot strategy (Asher et al., 2012; Sandoval et al., 2016), robot embodiment (Chaminade et al., 2012; Takahashi et al., 2014), payoff incentives (Hsieh et al., 2020), or group sizes in an intergroup competition (Fraune et al., 2019). Notably, some of these studies also recorded participants’ brain activity using functional magnetic resonance imaging (fMRI) while they played the game on-screen inside the scanner (Chaminade et al., 2012), sometimes after interacting with different robots outside the scanner (Takahashi et al., 2014). Nevertheless, the effect of robots’ communicative behavior during the game in real-time interaction with a physically present embodied robot has remained unexplored.

In this paper, we present a novel study of human decision-making under the influence of the communicative behavior (here, gaze) exhibited by a humanoid robot in an interactive setup. More precisely, we examine the influence of mutual versus averted gaze on participants’ behavior and neural activity, measured by electroencephalography (EEG), while they play the Chicken game against the iCub robot (Metta et al., 2010). We hypothesize that, as a key communicative signal, mutual gaze would affect participants’ performance and strategy in the decision-making task. Additionally, we hypothesize that the robot’s gaze would modulate participants’ brain responses during the decision period and after outcome presentation.

### Implementation of the Human-robot Chicken game

Participants sat in front of the iCub robot while the game was displayed on a screen placed horizontally on a table between them and the robot (Fig 1A). The experiment consisted of an adaptation of the Chicken game (Rapoport & Chammah, 1966). In the beginning of every round (hereafter “trial”), two car images, one per player, were shown on each side the screen. The cars started moving toward each other and stopped halfway before reaching the center. Then, the screen turned black and a fixation cross was shown for 1 second. After fixation crossed disappeared, the screen remained black for 5 seconds and both players had to choose their move between going straight or deviating. Critically, during this period, participants were instructed to look at the robot and the robot either look directly at the participant (hereafter labeled Mutual gaze) or avoid eye contact by looking to the side (hereafter labeled Averted gaze; see Fig 1B). At the end of the trial, the cars appeared on the screen again and displayed the action selected by the players. The possible outcomes were: i) both go straight and crash, resulting in the highest loss for both, ii) both deviate, obtaining the highest joint payoff, iii) only one goes straight obtaining the highest individual payoff. The points obtained (or lost) in the current trial were shown on each player’s side of the screen (see Fig 1A for the payoff matrix). Moreover, to maintain participants’ engagement, iCub verbally reacted to the outcome in 40 % of the trials within each block by randomly selecting one of the predefined utterances associated with that outcome (see Supp Table 1).

**Figure 1:**
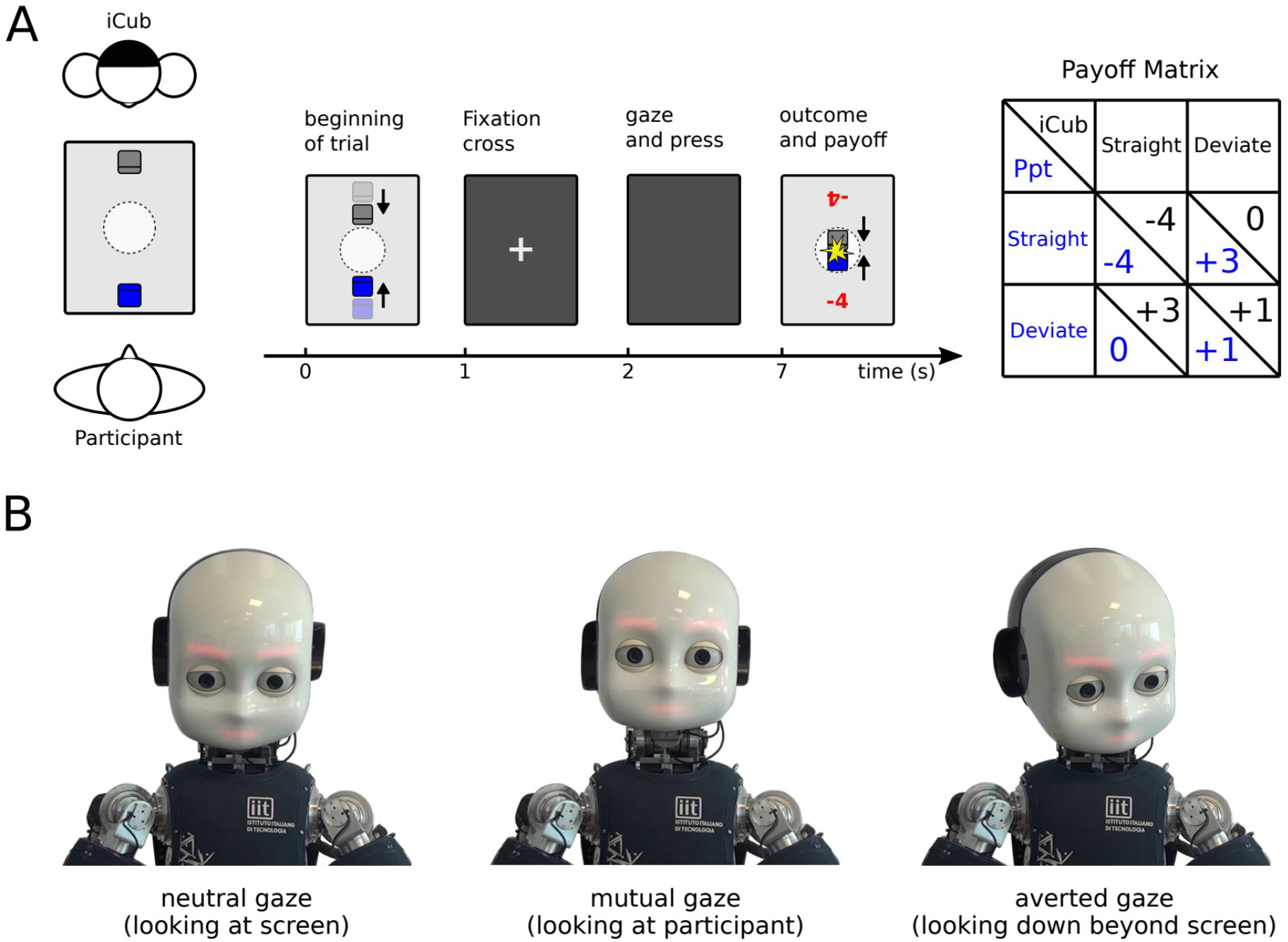
Human-robot Chicken game. **A)** Schematic illustration of the experiment, trial structure and payoff matrix. Participants sat facing the iCub robot while the game was displayed on screen placed between them. The players played an adaptation of the Chicken game in which we manipulated the robot gaze during the decision period (2-7 s). After looking at iCub, participants had to decide whether to go straight or deviate. The outcome of the trials and each player’s payoff was determined by the combination of the two players’ choices. **B)** During the entire trial, except for the decision period, iCub looked at the screen (neutral gaze). During the decision step, iCub either established or avoided eye contact with the participant (mutual or averted gaze, respectively)

Apart from the decision period when the robot’s gaze was manipulated (mutual or averted gaze), for the remaining time of the trial, the robot was looking at the screen, making periodic random saccades within that visual area. In this experiment, participants completed 250 trials divided in 5 blocks. They were assigned to either the group with 70% Mutual or 70% Averted conditions, meaning that in each block, iCub performed the type of gaze corresponding to the condition (e.g., mutual gaze in the 70% Mutual condition) in 70% of the trials within each block in a pseudo-randomized manner. The other type of gaze (e.g. averted gaze in the 70% Mutual condition) was performed in the remaining 30% of the trials. With this manipulation, we sought to investigate how participants’ behavior could be influenced by mutual gaze when established prior to decision-making (within-participants) and the effect of the degree of exposure to mutual versus averted gaze on their overall strategy in the game (between-participants).

Participants were told that their objective was to maximize their total score regardless of the robot’s. This was to make the always-go-straight strategy less appealing, because as much as it guarantees a higher score than the opponent, this strategy can lead to very low negative scores due to numerous crashes. Moreover, iCub followed a win-stay-lose-shift strategy (WSLS, Nowak & Sigmund, 1993) with a probability of 80%; meaning that it was most likely to repeat an action (i.e. stay) if it had led to a positive outcome in the previous trial and do the other action (i.e. shift) otherwise. Thereby, the robot’s sequence of choices had a certain structure (WSLS) that participants could capture without it being too obvious. Altogether, we gave both the incentive and the opportunity to participants to reason about the robot’s actions during the experiment.

## Results

### Performance in the human-robot Chicken game

Participants’ scores were markedly higher than the average score obtained when we simulated the risky always-go-straight strategy (Fig 2A). In fact, 45 % of the participants obtained a positive total score and 55 % obtained a higher score than iCub. However, total scores did not differ between conditions (70% Mutual versus 70% Averted, *t*(19)=0.50, *p*=0.61, *t*-test; Fig 2A). In addition, total scores were much lower than the average score obtained by simulation of the optimal strategy which consists in choosing the best action assuming that the opponent is using the WSLS strategy (Fig 2A). Indeed, a custom-made post-hoc questionnaire showed that 25 % of the participants thought that iCub had a strategy and only 12.5 % identified the actual WSLS strategy. These results are in line with those of pilot study we conducted prior to this experiment (see Materials and Methods and Supplementary Materials).

**Figure 2:**
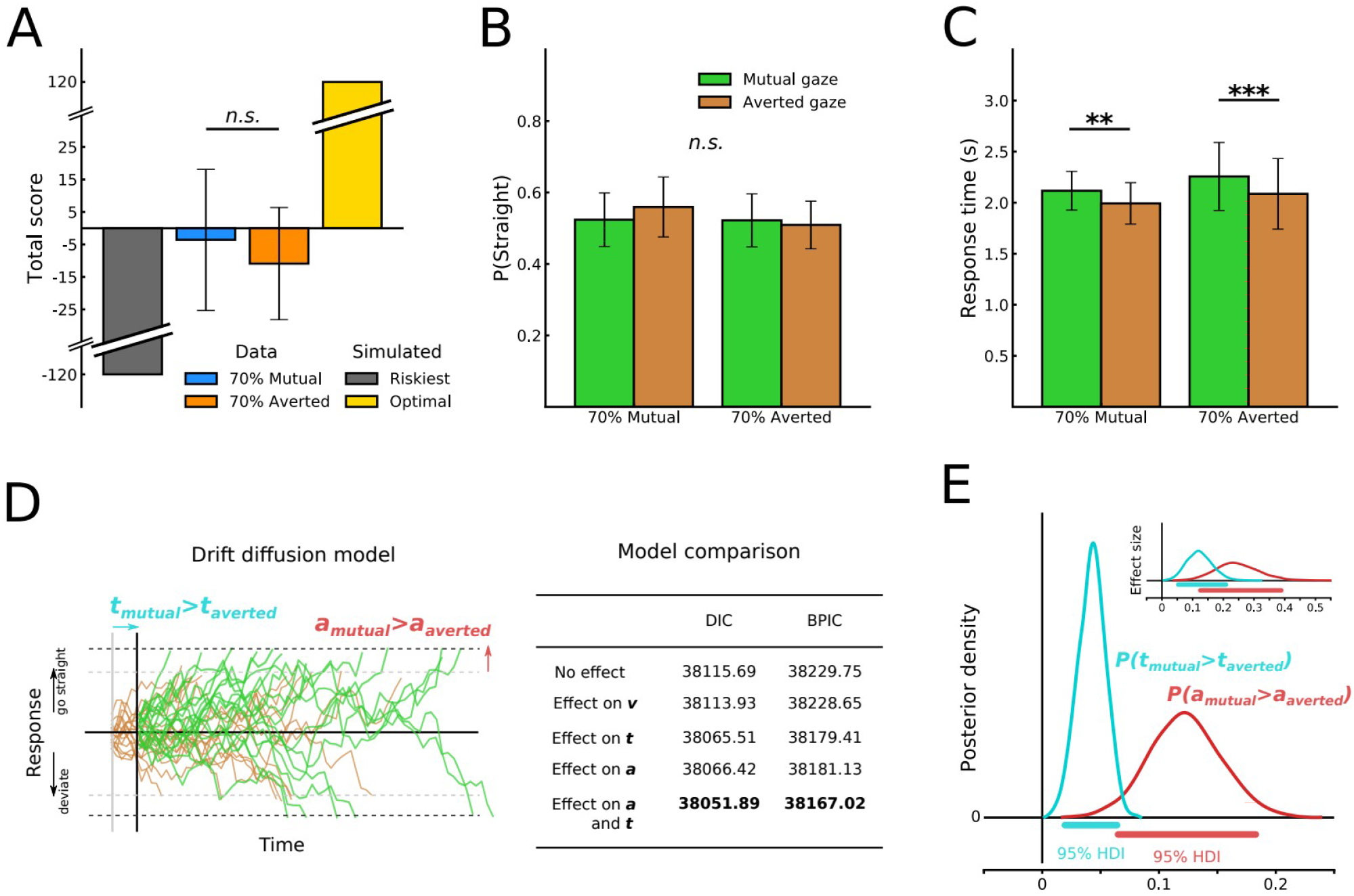
Participants’ performance and response times. **A)** Participants’ total scores did not differ between conditions. Average performance from simulations of two extreme strategies are provided for comparison (see text). **B)** No main effects or interaction of the condition or gaze type was found on the proportion of “straight” choices. **C)** The gaze type (within-subject) had a significant main effect on participants’ response times. **D)** The drift diffusion model describes decisions as a noisy drift process where an action is selected when the corresponding boundary is crossed. Five variants were tested assuming that the robot’s gaze had an effect of one, two or none of the model parameters. Upon Bayesian parameter estimation, the best fitting model was found to be the one imputing the difference in response times to an effect on both the non-decision time *t* and the decision threshold *a*. DIC: deviance information criterion. BPIC: Bayesian predictive information criterion. Lower values are better. **E)** Posterior density of the effect distributions for the best fitting model showing significant effects on both *a* and *t* parameters. HDI: highest density interval. Inset, effect sizes (see Materials and Methods). Error bars represent 95% confidence intervals. ** p < 0.01, *** p < 0.001. n.s., not significant at p > 0.05.

### Increased reaction times and decision threshold following mutual gaze

Using a two-way ANOVA with gaze condition within- and between-subject as factors, we found no significant main effect, all Fs<1, nor interaction, F=1.46, p=0.24, in the frequency of selected actions (Fig 2B). This means that the robot’s gaze did not influence participant’s subsequent choices, nor did the overall degree of exposure to the two gaze types. However, the same analysis showed a significant main effect of iCub’s gaze on participants’ mean response times, F=35.11, p<0.001, independent of the group they were assigned to (70% Mutual or 70% Averted). More specifically, participants responded faster following an averted gaze than following a mutual gaze (in 70% Mutual, M_averted_=1993 ms, SEM=106, M_mutual_=2117, SEM=99; in 70% Averted, M_averted_=2086, SEM=180, M_mutual_=2256, SEM=174; Fig 2C). This within-subject effect of gaze was also revealed by our pilot study where participants were exposed to the same amount of mutual and averted gaze instances (see Materials and Methods for procedure description and Supp Figure 1A for results). Thus, in this task, mutual gaze consistently elicited longer response times compared to averted gaze, whether the former was predominant, equally frequent or more occasional than the latter.

The delayed responses within-subjects following mutual gaze may suggest that direct gaze elicited more reasoning about iCub’s choices. To further investigate this hypothesis, we analyzed participants’ choices with the drift diffusion model (Ratcliff & McKoon, 2008). This widely used computational model assumes that decisions arise from relative evidence accumulation over time in favor of one of two alternatives (see illustration in Fig 2D). The main parameters include the drift rate *ν* (rate of evidence accumulation), decision boundary *a* (threshold to be reached to make a decision) and nondecision time *t* (dedicated to stimulus encoding and motor execution, for example). By fitting both alternatives’ selection rates and response time distributions with this model, we sought to infer the latent psychological processes explaining the delayed responses following mutual gaze. Specifically, a longer and more effortful reasoning process predicts an effect on decision threshold (Ratcliff & McKoon, 2008). Using hierarchical Bayesian parameter estimation (see Materials and Methods), we compared the fitness of different models assuming that the robot’s gaze had an effect on either the drift rate, the non-decision time or the decision boundary. Based on the deviance information criterion (DIC) and the Bayesian predictive information criterion (BPIC), we found that the two best fitting models were those presuming an effect on non-decision time or decision threshold: a difference in non-decision time explained the data slightly better in the main experiment (Fig 2D) whereas a difference in decision threshold explained the data better in the pilot study (Supp Fig 1C). Therefore, we also tested a model imputing the response delay to both non-decision time and decision threshold. This model fitted the data better than the others in both datasets (Fig 2D and Supp Fig 1C). In the main experiment, both the decision threshold *a* and the non-decision time *t* were significantly higher following mutual gaze compared to averted gaze (P(Δ*a*>0)=1.0, P(Δ*t*>0)=1.0) with a larger effect size for the former (Fig 2E). However, in the pilot study, only the difference in decision threshold reached significance (P(Δ*a*>0)=0.99, P(Δ*t*>0)=0.87; Supp Fig 1D). Taken together, these results show that the effect of gaze on response time is mainly driven by an effect of the decision threshold, which suggests a longer and more effortful decision process in case of mutual gaze, possibly due to the influence of more social components in this process.

### More synchronized alpha rhythm during mutual gaze

Behavioral analysis during decision period revealed a clear effect of iCub’s gaze on participants’ response time. We investigated whether this effect would also translate into distinct neural responses. For instance, time-frequency analysis can reveal different oscillatory activities across time in the human brain’s prominent rhythms that are linked to distinct neural processes (e.g., Ward, 2003). Therefore, we analyzed participants’ brain activity in the time period between the completion of the robot gaze movement (eyes and neck) and the average response time. We focused on theta and alpha bands which have been shown to be modulated by decision-making (Jacobs et al, 2006; Rajan et al, 2019), attention (Ward, 2003; Foxe & Snyder, 2011) and eye contact (Gale et al., 1972; Hietanen et al., 2008; Kompatsiari et al., 2019).

We did not observe any significant differences in theta frequency range. However, we found that participants responded with a higher alpha synchronization during mutual gaze compared to averted gaze in a parietal area (corrected for multiple comparisons p = .03; see Materials and Methods) during the time window 0.31-0.95 s relative to the initiation of robot’s head movement toward the gaze direction (see Figure 3). To locate the specific cluster of electrodes, we further averaged the data across the entire temporal period and compared the spatial data across conditions using the same analysis as above. Results showed that the parietal cluster consisted of six electrodes Pz, P3, P1, PO3, POz, P2. The averaged alpha values were lower in averted gaze M = 131.02 ± 4.34 compared to mutual gaze M = 323.97 ± 52.55. Increase in alpha power has been associated with brain processes related to the suppressing function of attention, i.e., suppressing cortical activity related to distractors or irrelevant information (Jensen, 2002, Cooper et al, 2003, for a review see Ward, 2003). Higher alpha synchronization in mutual gaze might indicate an increased need to suppress distraction related to the gaze exhibited by the robot, while focusing attention on task-related relevant information. The higher need for suppression of the irrelevant signal (iCub’s gaze directed at the participant) might have resulted in the observed behavior with longer reaction times for mutual, relative to averted gaze, trials.

**Figure 3:**
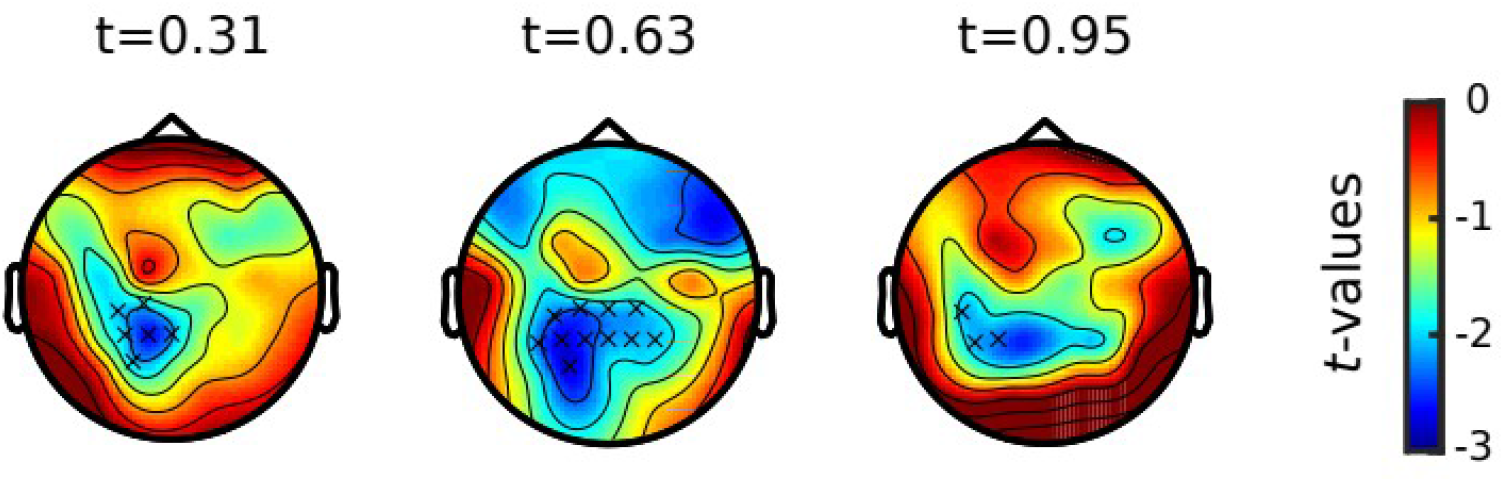
Scalp topographies between gaze types during decision period. Topographies show *t*-values maps of the difference in elicited alpha oscillations (8-12 Hz) between gaze conditions (averted gaze – mutual gaze). Statistically significant clusters are marked by x crosses. Differences between conditions were found by non-parametric cluster-based permutation tests. *t*-values are defined as the ratio of the difference between the estimated mean values of two conditions to its standard error. The topographies are depicted in a time range of t = 0.31 s to t = 0.95 s, relative to the establishment of the exhibited gaze (every 320 ms).

### Reasoning about the robot’s actions throughout the game

In order to assess how much participants reasoned about the robot’s choices during the game, we simulated three computational models taken from the literature (Hampton et al., 2008). These models make value-based decisions and compute different types of prediction errors to update the choices values over time (Fig 4A; see Materials and Methods for implementation details). The first model is a classical reinforcement learning model in which the action that was most rewarded in the recent past is more likely to be repeated (Rescorla and Wagner, 1972). The second model attempts to predict the opponent’s decision by estimating the probability that they select an action based on their recent history of choices. The model then selects the action which maximizes the gain (or minimizes the loss) based on the payoff matrix (Hampton et al., 2008). The third model builds on the second model and incorporates the influence of the player’s own recent choices on the opponent’s decisions by assuming that the other is tracking the player’s probabilities of choices too (Hampton et al., 2008). These models thus describe three levels of reasoning about the other (i.e. mentalizing). While the first model derives decisions solely from recent rewards (level 0), the second builds predictions of the opponent’s choices based on the recent history (level 1) and the third does so by also assuming a model of the other’s strategy (level 2).

**Figure 4:**
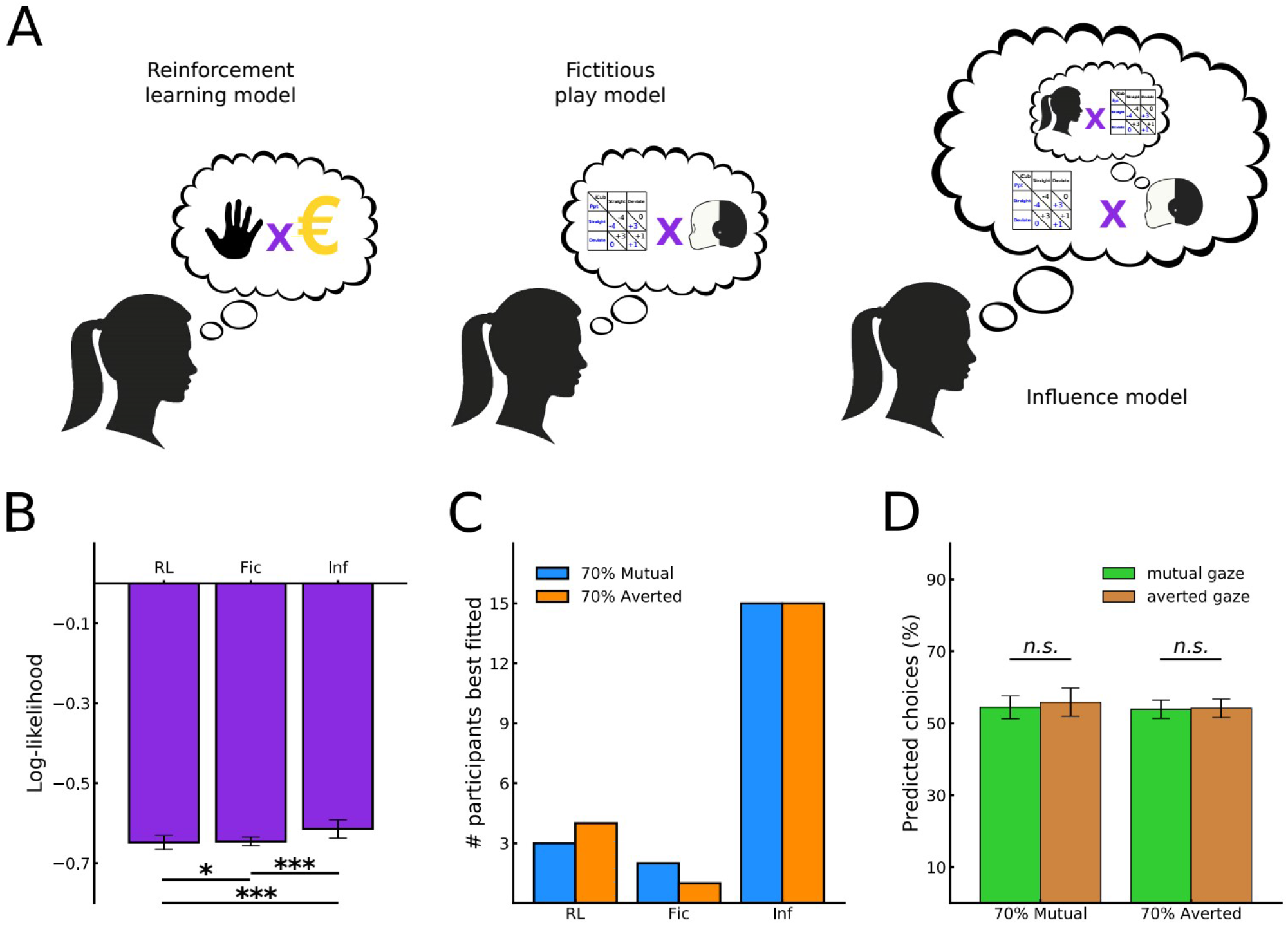
Computational models of participants’ degree of reasoning about iCub’s actions. **A)** Schematic illustration of the models. The reinforcement learning model (RL) makes decisions based on the recently selected actions and their outcome. The fictitious play model (Fic) makes decisions based on the game’s payoff matrix and the predicted action of the opponent. The influence model (Inf) does the same while assuming the opponent is also predicting the player’s choices and incorporating the influence of its own actions in its predictions of the opponent’s decisions. **B)** The overall log-likelihood of the influence model fitted to participants’ choices is significantly greater than the two other models, suggesting a high level of reasoning about iCub during the game. **C)** Considering the best fitting model for each participant individually did not reveal a strong difference between those exposed more to one type of gaze compared to the other. **D)** The percentage of trial-by-trial choices predicted by the model was similar following a mutual or averted gaze. Error bars represent 95% confidence intervals. * p < 0.05, *** p < 0.001. n.s., not significant at p > 0.05.

Upon optimization, we found that participants choice sequences altogether were best reproduced by the model with the highest level of mentalizing (log(L) across models, *H*=23.64, *p*<0.0001, Kruskal-Wallis test; Inf versus Fic, *U*=433.0, *p*=0.0002, Fic versus RL, *U*=547.0, *p*=0.007, Mann-Whitney tests; Fig 4B). By replicating previous findings in human-human strategic games (Hampton et al., 2008), we demonstrated that participants reasoned about iCub’s actions in a way that is similar to settings involving human opponents. Nevertheless, examining the best fitting model for each participant individually revealed little difference between our experimental conditions. In other words, participants exposed to 70% of mutual or averted gaze were fitted in similar proportions by the models (Fig 4C). Moreover, we examined the percentage of choices that were correctly predicted by the best fitting model for each participant. Results showed no significant effect of the trial’s gaze type within each condition (in 70% Mutual, *U*=171.5, *p*=0.22; in 70% Averted, *U*=192.0, *p*=0.41, Mann-Whitney test; Fig 4D). Overall, while suggesting that participants reasoned about the robot’s actions during the game, these computational models were unable to capture subtle differences which could further explain the effect of the mutual gaze on participants’ behavior.

### More self-oriented strategic patterns under higher exposure to averted gaze

To further investigate the influence of iCub’s gaze on participants’ decisions, we analyzed the frequency of occurrence of three patterns of strategic behavior in their choice sequences which could characterize the extent to which they incorporated information about their opponent in their decisions. The first pattern is ‘win-stay-lose-shift’ (WSLS), which is a typical decision-making strategy. It consists in repeating the previous action if it led to a positive outcome and abandoning it otherwise. This strategy is self-oriented, in the sense that it only requires information about the player’s own actions and outcome history and can be employed even in individual decision-making contexts. As a reminder, this was iCub’s strategy in 80% of the trials.

The second pattern that we examined is classical strategy in game theory called ‘tit-for-tat’ (T4T). The idea here is to reproduce the opponent’s previous choice. This strategy is known to be efficient in strategic games as it dissuades opponents from attacking and encourages them to cooperate by favoring actions which generate mutual gains. Additionally, we added another pattern labeled ‘stay-shift-imitation’ (SS-Imit). In this strategy, the player copies the opponent’s behavior in repeating the last trial’s choice or changing. In contrast to the first strategy, the two latter are other-oriented since they process information about the opponent’s history of actions and outcomes.

We analyzed participants’ choice sequences trial-by-trial looking for occurrences of these patterns of strategic behavior. Results showed that the pattern WSLS was found more frequently in the sequences of participants of the 70% Averted condition compared to 70% Mutual (*U*=108.5, *p*=0.006, Mann-Whitney test; Fig 5A). In other words, the occurrence rate of the self-oriented pattern was significantly lower under higher exposure to the mutual gaze. Interestingly, considering data from a pilot study, we found that when participants were equally exposed the both types of gaze, the occurrence rates of WSLS was similar to the 70% Mutual condition and also contrasted significantly with the 70% Averted condition (see Supplementary materials and Supp fig 3A). However, the difference between conditions for T4T and SS-Imit was not significant (T4T, *t(19)=-1.07, p=0.28*, SS-Imit, *t(19)=-1.02, p=0.30, t*-test; Fig 5A). Furthermore, we computed the average length of sequences of the strategic patterns to test for their recurrence over several trials. Again, we found a difference between the 70% Mutual and 70% Averted conditions for the WSLS pattern (*U*=120.0, *p*=0.015, Mann-Whitney test; Fig 5B) with longer sequences in the latter condition. On the other hand, the average length of sequences of the other patterns did not differ significantly between conditions (T4T, *U*=150.5, *p*=0.092, SS-Imit, *U*=189.0, *p*=0.38, Mann-Whitney test; Fig 5B). In summary, this analysis suggests that the likelihood of adopting self-oriented strategy was higher among those who were exposed mainly to averted gaze, relative to those who were exposed mainly or equally to mutual gaze.

**Figure 5:**
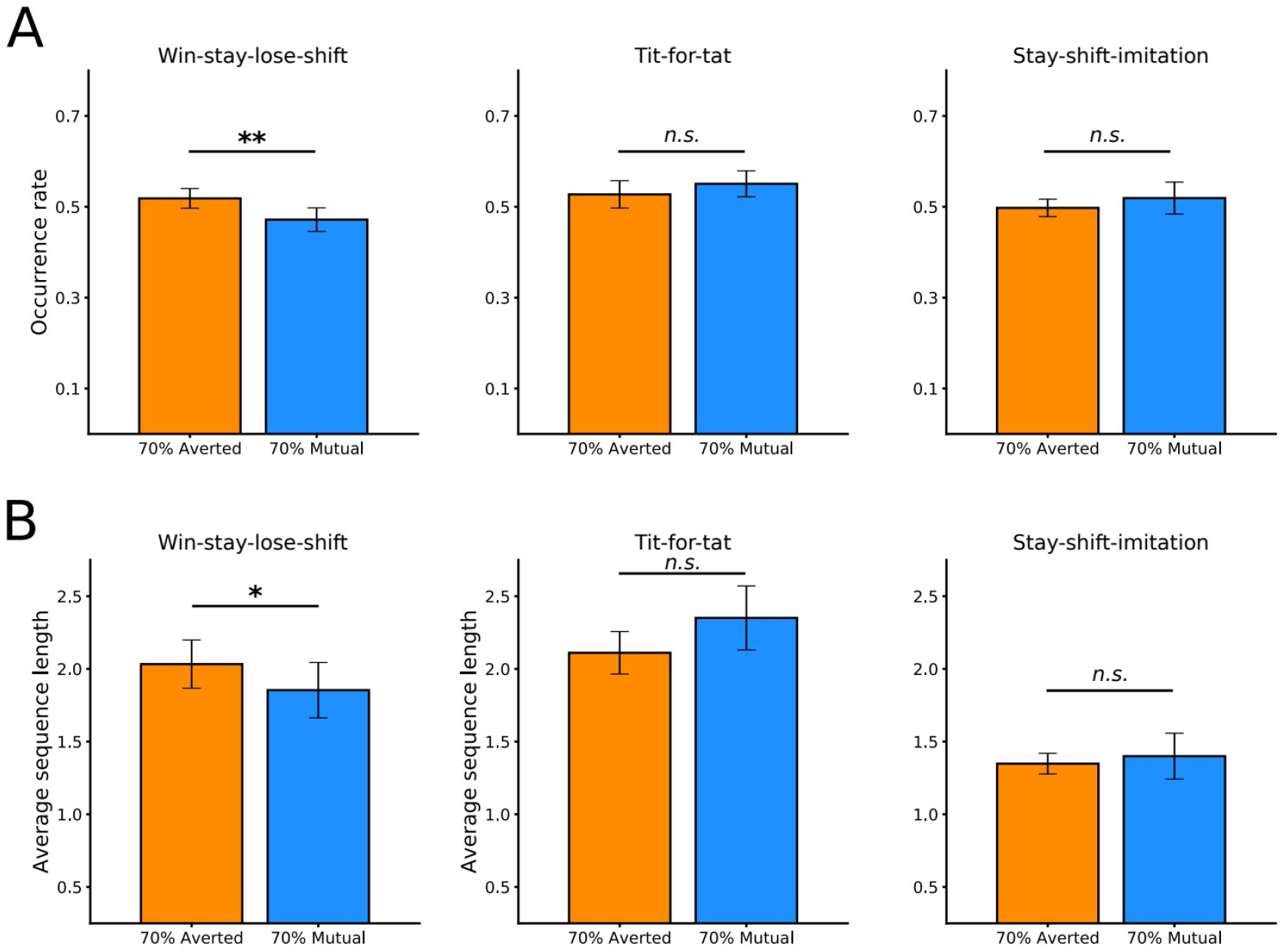
Patterns of self-oriented and other-oriented strategic behaviors in participants’ choice sequences. **A)** Occurrence rates of one self-oriented (win-stay-lose-shift, WSLS) and two other-oriented (tit-for-tat, T4T; stay-shift-imitation, SS-Imit) patterns (see text for detailed description). Self-oriented WSLS appeared significantly less in the choices of participant who were more exposed to the mutual gaze. No significant difference was found for T4T and SS-Imit. **B)** The average length of sequences of the WSLS pattern was significantly greater in the 70% Averted condition. No significant difference was found for T4T and SS-Imit. Error bars represent 95% confidence intervals. * p < 0.05, ** p < 0.01. n.s., not significant at p > 0.05.

### EEG markers of performance monitoring modulated by gaze

Because participants exhibited different strategic patterns depending on the degree of exposure to mutual or averted gaze, we reasoned that they would process their outcomes distinctly, as the analyzed strategies are inherently related to processing outcome and performance. At the neural level, performance and outcome monitoring are typically reflected in event-related potentials (ERPs) of the EEG signal, such as feedback-related negativity (FRN), error-related negativity (ERN) and the positive component P2 preceding FRN (e.g., San Martin, 2012, Gehring et al., 2013, Pollezi et al., 2008; Holroyd, & Coles 2002). Therefore, we examined ERPs during the period when participants had access to information about the outcome of the trial, i.e., within the broad time window from the offset of the black screen (the moment when the cars start moving after the action decisions have taken place) to the period after the score was presented (from 1000 ms to 3500 ms after onset of the animations; see Fig 1A). However, as our task was a dynamic game, with information about performance becoming available only gradually, the observed ERPs might not map onto classical ERP components reported in literature. Therefore, we refrained from typical component-based interpretations and focused on how activity evolved over time around maximum/minimum peak amplitude across consecutive 400-ms segments (see Materials and Methods for details about the procedure).

Our analyses showed that at first, exposure to mainly mutual or mainly averted gaze affected the EEG activity (Figure 6, T1, blue box), modulating the first positive peak of the entire sequence, with larger positivity M = 1.16, SEM = 0.27 for group with mostly averted gaze, compared to participants mostly exposed to mutual gaze, M = 0.43, SEM = 0.20, F (1, 37) = 4.44, p = .042, np^2^= 0.1. In a subsequent time window, there was no impact of gaze, but the EEG activity differed with respect to the outcome of participants’ decisions (Figure 6, T2, pink box). In particular, “winning” animations elicited a larger negative deflection, M=-1.01, SEM = 0.12 compared to the “losing” animations, M=-0.78, SEM = 0.1, F (1, 37) = 12.17, p = .001, np^2^= 0.25. This was then followed by another positive peak (Figure 6, T3, light green box) which also showed that outcome (but not gaze) modulated activity in this time period, with “winning” animations showing a larger positivity, M=0.92, SEM = 0.09 compared to the “losing” animations, M=0.6, SEM = 0.07, F (1, 37) = 17.19, p < .001, np^2^= 0.32. Last, in the segment following the presentation of the score (Fig 6, T4, yellow box), the positive peak was modulated both by gaze and feedback, but independently of each other. Participants mostly exposed to averted gaze showed a larger positive component M = 0.98, SEM = 0.21 compared to participants mostly exposed to mutual gaze, M = 0.48, SEM = 0.16, F (1, 37) = 4.6, p = .038, np^2^= 0.11. In parallel, winning feedback points showed a larger positivity, M=0.82, SEM = 0.13 compared to the losing feedback points, M=0.6, SEM = 0.15, F (1, 37) = 3.8, p =0.06, np^2^= 0.32.

**Figure 6:**
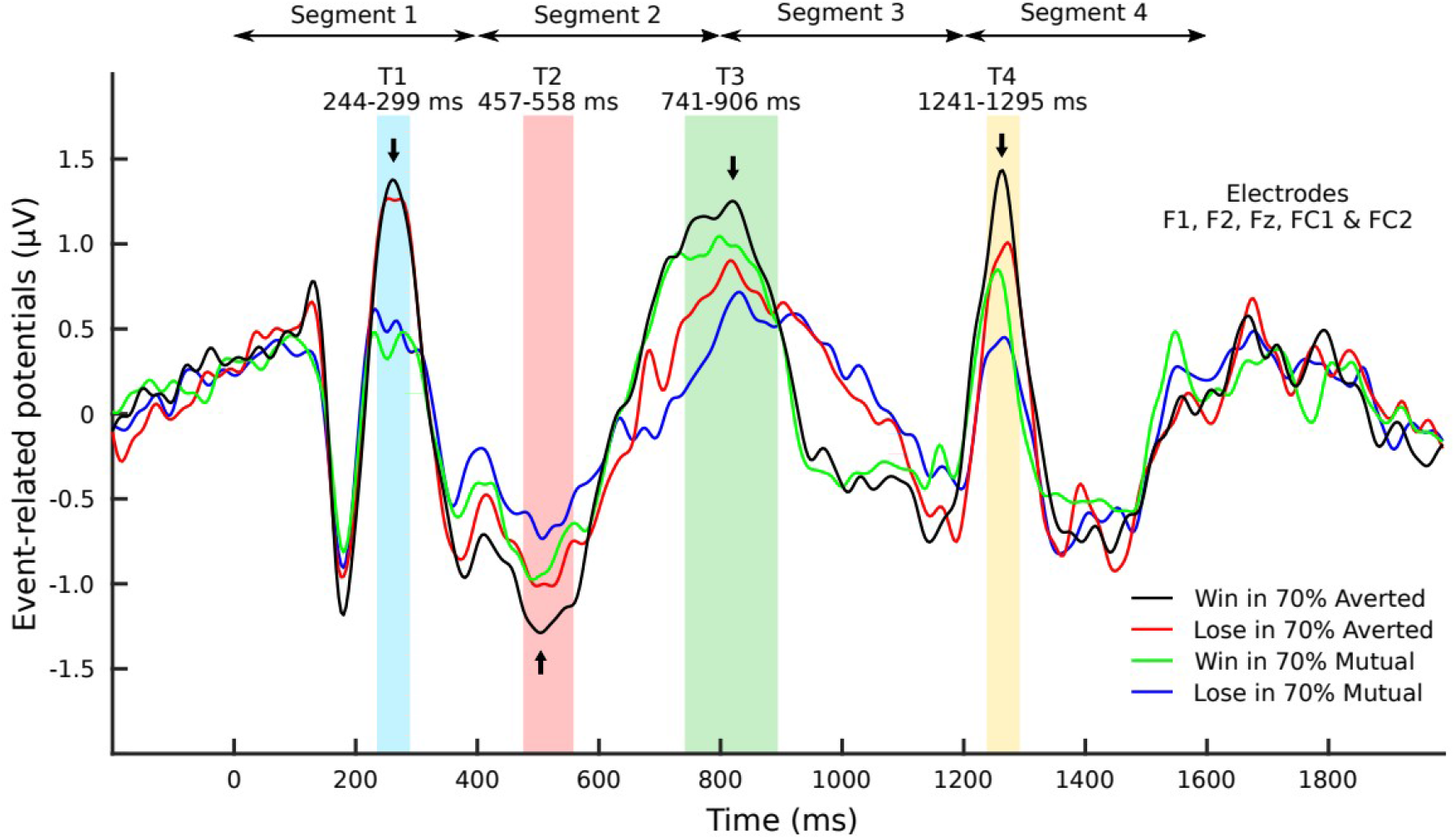
Grand-averaged ERP waveforms time-locked (t=0) to onset of feedback animation. The average is performed over F1, F2, Fz, CP1, CP2 electrodes on data from participants in the 70% Mutual and 70% Averted groups split into win (+1 or +3 points) or lose (0 or −4 points) trials. Activity over time around maximum/minimum peak amplitude across consecutive 400-ms segments showed differentiated responses to gaze condition in the first time window, then to outcomes in the following two time windows and finally to both gaze and outcomes. Division in segments: Segment 1: t = 0-400 ms; Segment 2: t = 400-800 ms; Segment 3: t = 800-1200 ms; Segment 4: t = 1200-1600 ms. Time windows of analyses (Tn) within each segment: Within Segment 1: T1 = 244-299 ms (light blue box) 10% around positive peak at 272 ms (black arrow); Within Segment 2: T2 = 457-558 ms (orange box), 10% around the negative peak at 508 ms (black arrow); Within Segment 3: T3 = 741-906 ms (green box), 10% around positive peak at 824 ms (black arrow); Within Segment 4: T4 = 1241-1295 ms (yellow box), 10% around the positive peak at 1268 ms (black arrow). Onset of score presentation: 1000 ms.

This pattern of results on how ERPs unfolded over the critical period of time, shows that the brain was initially affected by gaze: participants who were mainly exposed to avoiding gaze showed larger positivity. This is perhaps an after-effect of the preceding period where the two gaze types were displayed by the robot. However, this initial impact of gaze was in the later time windows overridden by the influence of the unfolding outcome of participants’ decisions, as participants observed the car animations and realized whether they have lost or won. In this time period, winning trials evoked a more pronounced deflection (larger positivity or negativity, dependent on whether the component was positive or negative), relative to losing trials. This clearly shows that at this point, the brain differentiated the wins from losses, and this was overriding the impact of gaze. Interestingly, though, upon receiving the scores, participants’ neural activity was affected both by feedback and by gaze condition. In other words, later stages of processing of performance (information about performance ultimately confirmed by the presented score) was modulated by the type of robot gaze to which participants were most exposed throughout the experiment.

## Discussion

Few studies have investigated the effect of social cues on human decision-making in naturalistic interactions. Our novel paradigm which made use of the humanoid robot iCub allowed us to overcome some of the methodological challenges of naturalistic interactive scenarios and enabled a real-time controlled interaction with a physically present and embodied agent. As such, it allows to shed light on the profound influence of others’ communicative signals (gaze in this case) on individuals’ behavior in strategic interactions.

In our experiment, participants played an adaptation of the Chicken game with iCub while the robot established or avoided eye contact with them prior to decision. Our results showed changes in participants’ behavior depending on iCub’s gaze. First, we observed that mutual gaze with the robot delayed participants’ responses. This difference in reaction times arose from a higher decision threshold in the mutual gaze condition compared to the averted gaze, thereby suggesting a longer and more effortful focus on the task. This behavioral effect was paralleled by a differential effect in synchronized alpha activity during the period of eye contact, with higher alpha synchronization compared to averted gaze. Alpha synchronization has been interpreted in literature as a marker of increased need for suppression of distractor information (for a review see Ward, 2003). In our experiment, iCub’s gaze was totally independent from its action and could thereby be considered by our participants as a distractor once the dissociation was detected. Therefore, mutual gaze might have required higher need for suppressing irrelevant signals in the environment (mutual gaze in this case), which presumably resulted in longer reaction times and higher decision threshold.

In addition to within-participants effects across different trials, we also analyzed strategic processes across the entire experiment, dependent on whether participants belonged to the group which was mostly exposed to the mutual gaze or to the avoiding gaze condition. We observed that while the type of gaze that participants mostly saw had no influence on choice frequencies or overall performance, patterns of strategic behaviors in participants’ choice sequence differed between conditions. Indeed, when exposed more to averted gaze, participants exhibited self-oriented patterns of choices more frequently. These results may suggest that higher exposure to averted gaze could make participants disengage more easily from the social interaction and rely more on self-centered information to make their decisions.

In search for the neural trace of performance/outcome monitoring processes, which would presumably underlie the different likelihood of adopting self-oriented strategy during the task, we examined ERPs of the EEG signal in the entire epoch when participants could monitor the outcome of their action decision. The results showed that the brain was indeed differentially processing information about winning and losing trials as this information was becoming available (as the car trajectories were gradually unfolding). Most importantly, being exposed mostly to mutual gaze or mostly to avoiding gaze affected the way the brain processed feedback. The group of participants who were exposed mostly to avoiding gaze showed larger amplitude than the group mostly exposed to mutual gaze. This differential neural processing might actually be one of the mechanisms underlying the behavioral result of differential strategies across the two groups of participants – those that were mostly exposed to averted gaze showed perhaps more sensitivity to feedback and, in result, showed larger proportion of self-oriented win-stay-lose-shift strategy relative to those that were mostly exposed to mutual gaze.

In summary, our results suggest that mutual gaze engages brain resources for managing social signals, which may or may not be relevant for the decisions to be made. In our experiment the robot gaze modulated response times, decision threshold, neural synchronization, as well as choice strategies and sensitivity to outcomes. Thus, two mechanical cameras were treated as a social signal, thereby providing striking evidence for the flexibility of human socio-cognitive mechanisms. As we build increasingly complex machines, it is therefore crucial that we endow them with adequate communicative behaviors.

## Materials and Methods

### Participants

43 participants were recruited for the main experiment, 3 were excluded due to high number of missing trials, excessive EEG artifacts, or technical issues during EEG recording. In total, data sets of 40 participants (mean age = 24.53 ± 4.5, 23 women, 5 left-handed) were analyzed for the main experiment, 18 participants (mean age = 30.38 ± 4.77, 10 women, 4 left-handed) for the first pilot study and 8 participants (mean age= 31 ± 4.87, 5 women) for the second pilot study. No statistical methods were used to predetermine sample sizes, but our sample sizes are comparable to similar previous studies (Wang et al., 2017, Kompatsiari et al., 2018). All participants were healthy and had normal or corrected-to normal vision. No participant who took part in the pilot study was recruited also for the main experiment, meaning that each participant took part only in one of the experiments. Participants were debriefed about the purpose of the study at the end of the experiment. The experiments were performed at the Istituto Italiano di Tecnologia (IIT). Participants who took part in the pilot studies were employed by IIT. Only external participants (participants of the main experiment) received honorarium (30 €) for their participation. All experiments were conducted in accordance with the ethical standards laid down in the 2013 Declaration of Helsinki and were approved by the local ethical committee (Comitato Etico Regione Liguria). All participants provided written informed consent prior to participation. Data were stored and analyzed anonymously. The data related to this study will be accessible online upon acceptance of the manuscript.

### Stimuli and apparatus

#### Task

The experiments were carried out in a noise-attenuated room. Participants were seated facing the iCub robot at opposite sides of a table at a distance of 130 cm. iCub is a humanoid robot (Metta et al., 2010), with 3 degrees of freedom in the eyes (common tilt, vergence, and version) and three additional degrees of freedom in the neck (roll, pitch, yaw). In our experiments, iCub was mounted on a supporting frame such that its eyes were at 124 cm from the floor which was estimated to be aligned with participants’ eyes. A 24-inch LCD screen was placed horizontally on the table such that both players (participant + iCub) could see the stimuli being displayed. The stimuli consisted of a series of animations describing the events occurring during the adapted Chicken game. In the beginning of every trial, two car images, one per player, were shown on each side the screen. The cars started moving toward each other and stopped halfway before reaching the center. Then, the screen turned black for 5 seconds and both players had to choose their next move between going straight or deviating. In the main experiment, a fixation cross was displayed prior to this step for 1 second. At the end of the trial, the cars appeared on the screen again and displayed the behavior selected by the players. Then the points obtained (or lost) in the current trial were shown on each player’s side of the screen. In 40 % of the trials within each block, iCub verbally reacted to the outcome by randomly selecting one of the predefined utterances associated with that outcome (see Supp table 1).

When the screen turned black (i.e. at the beginning of the decision step), iCub’s gaze changed to either make eye contact with the participant (mutual gaze) or avoid eye contact (averted gaze) by looking to the right or left of the table. During the rest of the trial, the robot was looking at the screen (neutral gaze), gazing at different fixation points randomly generated within the same field of view. Importantly, the target points to look at, in both the direct and the averted gaze conditions, have been selected in order to guarantee comparable conditions in terms of joints trajectories. During the rest of the trial, the robot was looking at the screen (neutral gaze), gazing at different fixation points randomly generated within the same field of view.

#### iCub behavior

The robot behavior was programmed using the YARP (Yet Another Robot Platform Python wrappers (Metta et al, 2006). We used iKinGazeCtrl, a 6-DOF gaze controller, to control iCub neck and eyes (Roncone et al., 2016). Specifically, Azimuth, Elevation and Vergence were provided to the controller in order to make iCub look at the desired 3D Cartesian coordinates (see Supp table 3). In addition, gaze shifts in head movements were embedded in order to make the gaze more naturalistic. The button press was controlled using a position controller, specifically YARP IPositionControl. The facial expression (always neutral in our experiment) and lip movements were controlled using the iCub faceExpressions application. Robot speech on the other hand simply consisted of audio files played on the main workstation with external speaker placed below the robot. The audio files were recorded from female human voices and edited using the free open-source Audacity software to increase the pitch. A custom-made software programmed in Python 3 and running on a with Ubuntu 20.04 LTS operating system was used to control iCub behavior, stimulus presentation, and data collection. Both players selected their actions by pressing switch buttons connected to a custom response box designed for converting input signals into regular keyboard key presses.

#### EEG apparatus

EEG was recorded from 64 electrode sites of an active electrode system using Ag-AgCl electrodes, at a sampling rate of 1 kHz (ActiCap, Brain Products, GmbH, Munich, Germany). FT9 and FT10 electrodes were displaced to F9 and F10 electrode positions to capture the horizontal ocular movements. All electrodes were referenced to FCz and re-referenced offline to average of all electrodes. Electrode impedances were kept below 10 kΩ throughout the experimental procedure.

### Procedure

Upon EEG head preparation and after receiving the task instructions, participants started a practice session to make sure that they were performing the task properly.

#### Pilot study 1

The experiment lasted approximately 45 min. Participants completed 200 trials divided in 20 blocks, 10 for each condition (direct gaze vs. averted gaze). There were breaks of 15 seconds between blocks. The order of the blocks was pseudo-randomized and it was counterbalanced across participants. Half of the participants experienced a sequence starting with a direct gaze block, while the other half experienced the opposite sequence. In each block, iCub performed the type of gaze corresponding to the current block (i.e. either direct or averted gaze) in 60% of the trials. To avoid a continuous exposure to one specific kind of gaze, the robot continued looking at the screen in the remaining 40 % of the trials. Here, the averted gaze was always directed to the right from the participant’s perspective.

In this pilot study, we collected also participants’ Galvanic Skin Response (GSR) in order to investigate whether it differed across conditions. However, due to the high number and variety of stimuli to which participants are exposed in this experiment, we found that the GSR signal was too noisy. We concluded that our experimental design is not compatible with this type of physiological response and discarded the GSR measure from the following experiments.

#### Pilot study 2

The procedure in this pilot study was exactly the same as in pilot study 1, except that the averted gaze was directed to the left from the participant’s perspective. The reason was that in Pilot study 1, both the robot’s and the participant’s button were placed on the right side of the table from the participant’s perspective. We thus wanted to test whether the difference in response time observed in Pilot Study 1 was due to a higher action readiness caused by the averted gaze being directed to the side on which the buttons were placed. Upon verifying that the pattern of results in Pilot Study 2 was in line with Pilot Study 1, and thus the pattern was not related to the direction of the robot gaze (direction to the right), we did not further analyze this dataset. Results related to the “Pilot study” in the main text and supplementary material only concern data collected in Pilot study 1.

#### Main experiment

The experiment lasted approximately 55 min. Participants completed 250 trials divided in 5 blocks. Between-block breaks lasted 90 seconds after the first, second and fourth block, whereas participants could take a longer break after the third block and decide when they wanted to resume the experiment. This experiment consisted of two between-participants conditions: 70% Mutual (20 participants) or 70% Averted (20 participants). For example, participants in the 70% Mutual condition experienced the mutual gaze in 70% of the trials and the averted gaze in the remaining 30% of each block. The sequence of trials was pseudo-randomized and participants in the 70% Averted condition experienced the exact opposite sequence. The direction of the averted gaze (left or right) was counterbalanced across two subgroups in each condition. Put differently, participants were assigned to one of the following four subgroups: 70% Mutual with averted gaze to the right, 70% Mutual with averted gaze to the left, 70% Averted with averted gaze to the right, 70% Averted with averted gaze to the left.

Prior to the main experiment, we collected resting state EEG data with eyes closed and eyes open. After all experiments were over, participants were also asked to answer two questionnaires: one custom-made questionnaire related to self and robot’s strategy, and a competitiveness index questionnaire (Houston et al. 2000). Informal discussions after the experiment confirmed that all participants noticed the gaze manipulation. The resting state and the competitiveness index questionnaire were aimed at targeting possible individual differences, which is out of the scope of this paper. Therefore, these data are not reported here.

### Drift diffusion model

The drift diffusion model is a well-established computational model which describes the intra-trial decision process in two-alternative forced choice tasks as the relative accumulation of evidence over time (Ratcliff & McKoon, 2008). The main parameters include the drift rate *ν* (rate of evidence accumulation), decision boundary *a* (threshold to be reached to make a decision) and non-decision time *t* (dedicated to stimulus encoding or motor execution, for example). Choices are represented by one of the boundaries. A noisy drift-process accumulates evidence until it crosses one of the two boundaries which indicates the selection of the associated alternative. Described as such, the model aims to infer the latent psychological processes underlying subjects’ decisions by fitting the two alternatives’ selection rates and response time distributions.

To fit the model to our data, we used the HDDM (hierarchical drift diffusion model) open-source python package (Wiecki et al., 2013). HDDM relies Markov chain Monte Carlo (MCMC) algorithm to perform Bayesian estimation of the model’s parameters such that subject parameters are assumed to be drawn from the group distribution (see Kruschke (2013) for details about Bayesian parameter estimation in a non-hierarchical setting). The toolbox also allows flexible model construction where multiple posterior distributions are estimated for specific parameters which are assumed to be affected by certain experimental conditions. To evaluate within-subject effects (e.g. influence of gaze type in our experiment), individual parameters can be described by a linear model with independent variables specified as covariates.

For each one of the five model variants that we tested (see text and Fig 2D), we set the MCMC sampling algorithm to draw 30000 samples from which the first half was discarded as burn-in. Then MCMC chain convergence was first assessed by visually inspecting the trace, the autocorrelation, and the marginal posterior for each parameter chain (Wiecki et al., 2013). In addition, by running 3 independent parameter estimation processes for each model, we also measured the Gelman-Rubin convergence criterion (Gelman and Rubin, 1992). The value of this statistics was smaller than 1.02 for all the chains, which indicates good convergence for all the models. Once the convergence checked, we performed model comparison based on the deviance information criterion (DIC) and the Bayesian predictive information criterion (BPIC). Lower values of these two criteria indicated better fitness. We then inspected the posterior distributions to assess the extent to which the models accounted for the effect of the experimental manipulation. Specifically, by defining a linear model to describe the effect, the fitting procedure estimates a posterior distribution of the difference *Δ* between the two conditions. The size of the portion of this effect distribution that is above zero was used to determine the significance of the effect. Similarly to Kruschke (2013), effect sizes were estimated as 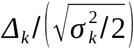 where *Δ_k_* is the estimated difference between conditions for a parameter *k* and σ_k_ is the estimated standard deviation.

### Value-based decision-making models

This family of computational models aims to describe the inter-trial learning process underlying decision-making. In this framework, choices are made based on the available options’ values, which are updated trial by trial. In this paper, we used this type of models to study the mechanisms on which participants relied during the game. Specifically, the degree to which participants reasoned about the robot’s action. To do so, we readapted three computational models taken from the literature (Rescorla and Wagner, 1972; Hampton et al., 2008). Model 1 is a reinforcement learning where most recently rewarded actions are selected and involves no reasoning about the opponent. Model 2 estimates the probabilities of opponent’s actions based on recently selected choices then constructs its action values by combining the estimated probabilities with expected outcomes from the payoff matrix. Model 3 adds another level of reasoning about the opponent by integrating the influence of its own recent choices on the opponent’s decisions. These models are detailed in Supplementary materials.

We optimized the models’ hyperparameters to fit each participant’s sequence of choices by maximizing the log-likelihood:

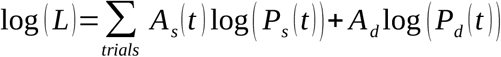

The optimization procedure was performed using the fmin function from the scipy.optimize Python module (version 1.5.0) by minimizing the negative log likelihood −log(L). To avoid local minima, we repeated the procedure with n initial points (n = 10 times the number of model hyperparameter).

Prior to fitting the actual data from our participants, we sought to assess the model fitting procedure. To do so, we simulated 100 surrogate subjects with each model, then fitted all models to all simulated data. As a result, we found that all models had good recovery rates (Supp fig 2A), though Model 2 (Fic) had a higher confusion rate compared to the reinforcement learning and the influence models. This indicated that this model was less likely to emerge as the best fit for participants’ data in the actual fitting procedure.

Then, upon optimization of the models’ hyperparameters to fit the experimental data, we performed an additional validation step. We simulated again the best model of each participant with the best fitting parameters in order to check that the observed behavior was properly reproduced. Specifically, we averaged the probability of going straight, the probability of repeating the same choice as the previous trial, and the total score over 10 repetitions for each best fitting model and compared the simulated performance to the actual participants’ performance. Linear regressions confirmed that simulated and experimental data were strongly correlated (r > 0.75) for the first two measures and moderately correlated (r > 0.5) for the third measure (Supp fig 2B, 2C and 2D).

### EEG data analysis

#### Preprocessing

EEG data were analyzed using MATLAB® version R2018b (The Mathworks Inc., 2018), the EEGLAB (Delorme & Makeig, 2004), FieldTrip toolboxes (Oostenveld et al.,2011) and customized scripts. The data were down-sampled to 250 Hz, while a band-pass filter (0.5–100 Hz) and a notch filter (50 Hz) were applied. The signal was re-referenced to the common average of all electrodes. Data from one participant (from the 70% Averted group) were excluded only from EEG analyses due to a large number of noisy electrodes (> 10), resulting in N=39 for EEG analyses.

#### Time-frequency analysis during decision (and gaze) period

To analyze participants’ neural activity during the decision period, data were segmented into trials of 6 s length, including the fixation cross (1 s) and the gaze plus response period (5 s). Each trial was baseline-corrected by removing the values averaged over a period of 500 ms during the fixation cross (0.2 to 0.7 s relative to stimulus onset). The specific baseline window was chosen in order to ensure a relatively large task-unrelated window. First, the extracted segments were visually inspected and trials with large artifacts (i.e., muscle movements and electric artefacts) were removed. The same procedure was applied to the removal of bad channels. The mean number of the remaining trials was similar across gaze conditions (mutual or averted) and equal to 159.87 ± 9.25 (out of 175) for trials of the main gaze (e.g. mutual gaze trials for the 70% Mutual group) trials and 69.77 ± 4.14 (out of 75) for the other trials on average. On average, 3.28 ± 1.43 electrodes were removed and interpolated afterwards. The remaining artefacts, i.e., muscular activity, ocular activity and channel noise were removed by applying independent component analysis (ICA). The number of removed ICA components was equivalent to 26 ± 8 on average. After the artefact removal, noisy channels were spatially interpolated.

Time-frequency representations (TFRs) of oscillatory power changes were computed separately for each condition (mutual gaze, averted gaze) for the abovementioned period. Time-frequency power spectra were estimated using Morlet wavelet analysis based on varying cycles to allow for high spectral resolution in lower frequencies (3.5 cycles at the lowest considered frequency: 2 Hz) and high temporal resolution for higher frequencies (18 cycles at the highest considered frequency: 60 Hz). Time steps were set to 10 ms while frequency steps were set to 1 Hz (Oostenveld et al., 2011). All trials were averaged for each condition across participants and the butterfly plots were used to inspect for potential ERPs during the period of interest. ERPs were observed only in the baseline correction period and the evoked activity was removed. Subsequently, an absolute baseline correction for each trial was performed by subtracting the average induced oscillatory activity of the 0.2 to 0.7 s during the fixation cross period (Premoli et al., 2017). This baseline correction was used to avoid any task-related timefrequency activity. Subsequently, TFRs were averaged across trials per experimental condition. The epoch of interest consisted of actual mutual/averted gaze phase before participants’ average response time. For this reason, as initial time point of statistical analysis we annotated the establishment of mutual/averted gaze; and as the final time point, the participants’ average response time. TFRs were cropped to the phase of interest.

Data were averaged to calculate power within theta (4-7) and alpha (8-12) frequency bands. In each frequency range, spatio-temporal data across conditions were compared by performing a nonparametric cluster-based permutation analyses (using a Monte-Carlo method based on paired t-statistics) (Maris & Oostenveld, 2007). Samples between gaze conditions with a t-value exceeding an a priori threshold of p < .05 were clustered on a temporal and spatial adjacency basis. Subsequently, comparisons were performed for the maximum values of summed t-values. A permutation test (i.e., randomizing data across conditions and re-running the statistical test for 1500 times) was used to approximate a reference distribution of the maximum of summed cluster-level t-values. Clusters consisting of a minimum of two electrodes were considered statistically significant at an alpha level of .05 if < 5% of the permutations used to construct the reference distribution yielded a maximum cluster-level statistic larger than the cluster-level value observed in the original data.

#### Performance monitoring ERPs

In order to examine how information about participants’ performance in our task was processed at the neural level, we focused on the event-related potentials ERPs during the period when participants could already understand their performance within a given trial, i.e., within the broad time window from the offset of the black screen (the moment when the cars start moving after the action decisions have taken place) to the period after the score was presented (score was presented 1000 ms after onset of the animations), cf. Fig 1A. Visual inspection of the signal (cf. Fig. 6) showed clear ERP components. However, the design was not tailored towards specific ERP components. Our task was a dynamic game, with the information about performance becoming available only gradually, so the observed ERPs might not map onto more classical ERP components reported in literature. Therefore, we decided to refrain from classical component-based analyses or interpretations, and we focused rather on how activity evolved over time from the moment of onset of animations. We analyzed the data focusing on the average activity around maximum/minimum peak amplitude across consecutive 400-ms segments. As we refrain the interpretation of the components in terms of classical ERP components, we simply refer to our analyses in terms of min/max peak amplitude within various time windows (i.e. rather than naming the ERPs as, for example, P1 or N1 components).

Data were segmented into trials of 2.2 s length, including 200 ms before the feedback animation, the feedback animation (t = 0-1 s) and the first second of score presentation (t =1-2 s). Each trial was baseline-corrected by removing the values averaged over a period of 200 ms before the feedback animation. After preprocessing, the average remaining trials was similar across groups and equal to: win trials: 117.15 ± 9.0, lose trials: 118.61 ± 8.21, the number of interpolated electrodes was equal to: 3.28 ± 1.43 (same electrodes with TFA analysis), and the number of removed ICA components was equal to: 18.66 ± 5.88. Event-related potentials were calculated and low-pass filtered to 30 Hz. We calculated the minimum and the maximum voltage values within segments of 400 ms starting from the onset of the feedback animation and until 600 ms after score presentation. We focused on a set of frontocentral electrodes, i.e., F1, F2, Fz, FC1, FC2, that are sensitive to feedback presentation (San Martin, 2012; Gehring et al., 2013; Pollezi et al., 2008). For every maximum/minimum voltage peak in the respective time window, we computed the average activity over a window of ±10% relative to the peak. The computed average amplitude values were submitted to a repeated-measures ANOVA with feedback type (win, lose) as a within-subjects factor, and gaze group (70% mutual gaze, 70% averted gaze) as a between-subjects factor.

### Statistical analysis

Details about the statistical tests were reported in the main text. Error bars in figures indicate 95% confidence intervals. For one-factor analyses, parametric statistical tests were used when data followed a normal distribution (Shapiro test with p > 0.05) and non-parametric tests when they did not. As parametric tests, we used t-test when comparing two groups or ANOVA when more. As nonparametric tests, we used Mann-Whitney test when comparing two independent groups, Wilcoxon test when comparing two paired groups and Kruskal-Wallis test when comparing more than two groups. These tests were applied using the scipy.stats Python module. They were all two-sided except Mann-Whitney. Two-factor analyses were performed using the JASP software. In all statistical tests, p > 0.05 was considered to be statistically non-significant.

## Acknowledgments

The authors would like to thank Abdulaziz Abubshait, Francesco Bossi, Francesca Ciardo, Jairo Perez-Osorio and Cesco Willemse for their advice and feedback, Adam Wojciech Łukomski for his help in developing the experiment software, and Marco Crepaldi for his work on the response box controller. This project has received funding from the European Research Council (ERC) under the European Union’s Horizon 2020 research and innovation program (grant awarded to A.W., titled “InStance: Intentional stance for social attunement.” G.A. no.: ERC-2016-StG-715058). The content of this paper is the sole responsibility of the authors. The European Commission or its services cannot be held responsible for any use that may be made of the information it contains.

## Author contributions

M.B., K.K. and A.W. designed the study. M.B. and D.D.T. programmed the experiment. M.B., K.K. and I.Z. collected the data. M.B. and K.K. analyzed the data. M.B., K.K. and A.W. wrote the manuscript.

## Supplementary material

### Results from the pilot study

In the pilot study, participants completed 200 trials divided in 20 blocks, 10 for each condition (mutual gaze vs. averted gaze). The order of the blocks was pseudo-randomized and it was counterbalanced across participants. Half of the participants experienced a sequence starting with a mutual gaze block, while the other half experienced the opposite sequence. In each block, iCub performed the type of gaze corresponding to the current block (i.e. either mutual or averted gaze) in 60% of the trials. To avoid a continuous exposure to one specific kind of gaze, the robot continued looking at the screen in the remaining 40 % of the trials (see Figure 1B).

Results showed that 47.1 % of the participants obtained a negative total score, whereas 58.82% obtained a higher score than iCub. In addition, a custom-made post-hoc questionnaire showed that 38.88 % of the participants thought that iCub had a strategy, 11 % identifying the actual win-stay-lose-shift strategy. We found no significant difference in the frequency of selected actions (straight versus deviate, Z= −.64, p = .52, Wilcoxon signed-rank test; see Supplementary Figure 1A). However, mean response times differed significantly between gaze conditions (Z= −2.86, p = .004, Wilcoxon signed-rank test; see Supplementary Figure 1B). More specifically, the mutual gaze elicited longer response times compared to the averted gaze (M_direct_=1879.96 ms, SEM=169.38; M_averted_=1747.15 ms, SEM=181.42; Fig 1B).

We also analyzed participants’ choices using hierarchical Bayesian estimation of the parameters of the drift diffusion model (Ratcliff & McKoon, 2008). We found that the model assuming an effect of the gaze on the decision threshold parameter *a* explained the data better than those assuming an effect on the drift rate *v* or non-decision time *t* (lower DIC and BPIC values; see Supplementary Figure 1C). Given the results of the same analysis for the main experiment (see Figure 2C), we also tested the model assuming an effect on both *a* and *t* and found that it fitted better than the previous models. However, while the effect on *a* showed strong significance and large effect size (P(Δ*a*>0) = 0.99; see Supplementary Figure 1D), the effect on *t* did not reach significance (P(Δ*t*>0) = 0.87; see Supplementary Figure 1D; see Materials and methods of the main text for details about Bayesian estimation of the drift diffusion model parameters)

### Supplementary results on patterns of strategic behaviors in the pilot study

In contrast to the main experiment, the pilot study included an equal percentage of mutual and averted gaze for each participant. Therefore, this data can offer a complementary view on the effect of different degree of exposure to each type of gaze. We found that the occurrence rates of win-stay-lose-shift (WSLS) in the data of pilot experiment were similar to those of the 70% Mutual condition in the main experiment (see Supplementary Figure 3A). In other words, participants who were the least exposed to the mutual gaze (i.e. the 70% Averted condition) showed a significantly higher rate of occurrence of the self-oriented pattern (WSLS) than those who were exposed to equal (pilot data) or higher number of mutual gaze.

In addition the patterns reported in the main text, we analyzed a fourth pattern labeled ‘counter-win-stay-lose-shift’ (C-WSLS). Given that in our experiment iCub mostly followed the ‘win-stay-lose-shift’ strategy, the optimal strategy for a player aware of iCub’s strategy is to counter with the action which maximizes the gain or minimizes the loss depending on the predicted robot action. For instance, if iCub went straight and lost in the previous trial, it is likely to deviate in the following trial, in which case going straight brings the highest outcome. However, closer examination of the WSLS and C-WSLS strategies in the context of this experiment revealed that they systematically select opposite actions (see Supp table 2), thus rendering their comparison in terms of occurrence rate rather trivial. Nevertheless, the length of sequences of these two strategic patterns indicating their recurrence over several trials showed distinct results: sequences of WSLS were longer in 70% Averted relative to 70% Mutual while no significant difference was found for C-WSLS.

### Detailed description of the value-based decision-making models

Similarly to Hampton et al. (2008), we used three computational models with different levels of reasoning about the opponent’s actions in order to analyze participants’ strategies. All three models rely on a form of prediction error to estimate the value of each action (see details below). The agent then decides which action to perform (go straight *s* or deviate *d*) based on the logistic sigmoid function *f*. For example, the probability *P_s_* of going straight at trial t is:

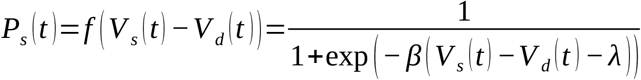

where *V_a_* is the value of action *a*, *λ* is a parameter describing a bias toward going straight, and *β* is the inverse temperature parameter which determines the degree of stochasticity in the agent’s choice (high values increase the propensity of choosing the action with the highest value while low values reduce the difference between action probabilities). The bias parameter was found to improve model fitting and recovery.

#### Level 0 – Reinforcement learning model

This is a classical model using the Rescorla-Wagner rule. Agents learn the value of each action based on the outcome following the performance of these actions in the past. After each trial *t*, the value *V_a_* of previous action *a* is updated as follows:

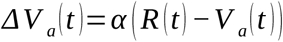

where *R* is the reward experienced at trial *t* and *α* the learning rate. Thus, this model tracks the action values using the prediction error determined by the difference between the expected and the experienced reward. Only the value of the selected action is updated after each trial and the values are plugged in the decision rule defined above.

#### Level 1 – Fictitious play model

This model introduces a form of reasoning about the other where the agent estimates the probability *P*^*Opp*^_a_ that the opponent chooses a certain action as follows:

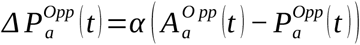

where *A^Opp^_a_* is a binary variable which equals 1 if the action *a* is performed by the opponent at trial *t*, and *α* is the learning rate. The opponent then attributes values to actions based on the game’s payoff matrix and the opponent’s probability of actions:

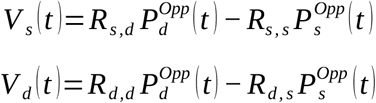

where *R_i,j_* is the outcome obtained when the agent performs action *i* and the opponent performs *j* according to the payoff matrix. Thus, this model tracks the probability of the opponent’s actions using the prediction error determined by the difference between the expected probability of action and the observed realization of an action.

Considering that *P^Opp^_d_* = 1 - *P^Opp^_s_* and given the payoff matrix used in our experiment, the decision rule can be expressed as follows:

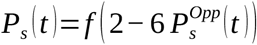

#### Level 2 – Influence model

This model builds on the fictitious play model to take into account the influence of the agent’s actions on the opponent’s choices. This is done by assuming that the latter is using a fictitious play strategy. The estimated probability of the opponent’s action is then computed as follows:

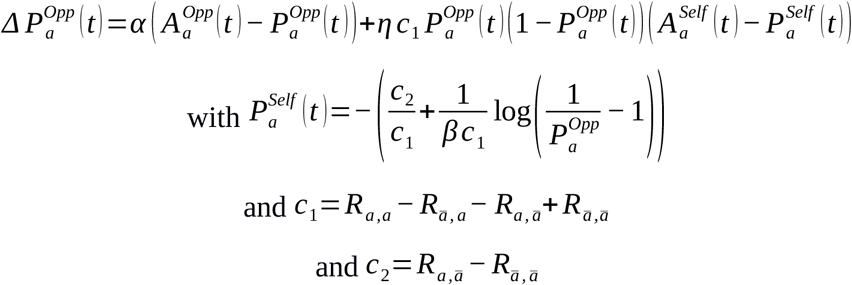

where *A^S^_a_* is a binary variable which equals 1 if the action a is performed by the agent at trial *t, η* is the influence rate, *c_1_* and *c_2_* are constants computed from the payoff matrix. For example, to estimate *P^Opp^_s_*, we calculated that *c_1_*=-6 and *c_2_*=2. Additionally, *P^Self^_a_* is a second-order estimation of the opponent’s estimation of the probability that the agent performs action *a*. This added influence term is approximated by plugging the update rule of the probability estimation in the equation for value calculation. The second-order probability estimation is obtained by inverting the decision rule (see Hampton et al., 2008). This model thus incorporates an additional form of prediction error determined by the difference between the prediction of the expected probability of the player’s own action as estimated by the opponent and the observed realization of the action.

To reduce the number of hyperparameters in this model, we also tested a variant in which *η*=*α*. This variant was found to have similar results compared to the full version and was thus preferred for its simplicity. Thereby, all three models (RL, Fic and Inf) have the same number of hyperparameters (=3): *α, λ* and *β* which were constrained between 0 and 1, 0 and 1 and 0 and 10 respectively. In the optimization procedure, they were randomly initialized by sampling the following distributions: *α* ~ *Beta* (1.1,1.5), *λ* ~ *Beta* (1.1,1.1), *β* ~ *Beta* (1.1,1.5,10). These distributions were defined using the Python scipy.stats package and when indicated, the third argument described the scaling parameter. The fitting procedure is explained in the Materials and methods sections of the main text.

**Supplementary Figure 1:**
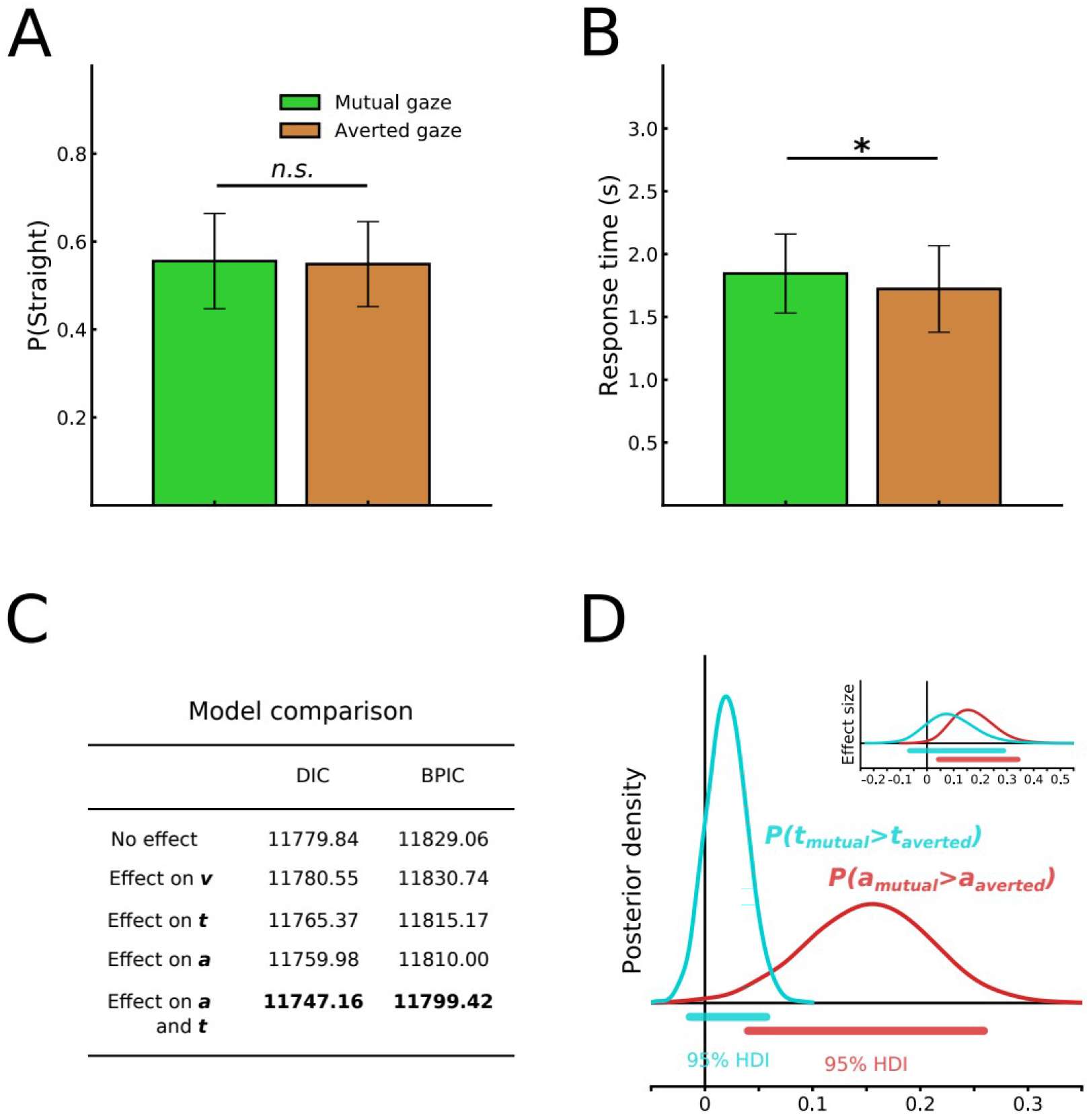
Participants performance and response time in the pilot study. **A)** No significant effect of the gaze was found on the proportion of “straight” choices. **C)** Response time was significantly longer following mutual gaze compared to averted gaze. **D)** Model comparison for five variants of the drift diffusion model assuming that the robot’s gaze had an effect of one, two or none of the model parameters. The best fitting model was found to be the one imputing the difference in response times to an effect on both the non-decision time t and the decision threshold a. DIC: deviance information criterion. BPIC: Bayesian predictive information criterion. Lower values are better. **E)** Posterior density of the effect distributions for the best fitting model showing a significant effect on decision threshold a but not on non-decision time t. HDI: highest density interval. Inset, effect sizes (see Materials and methods). N=18 in A and B. Error bars represent 95% confidence intervals. * p < 0.05. n.s., not significant at p > 0.05.

**Supplementary Figure 2:**
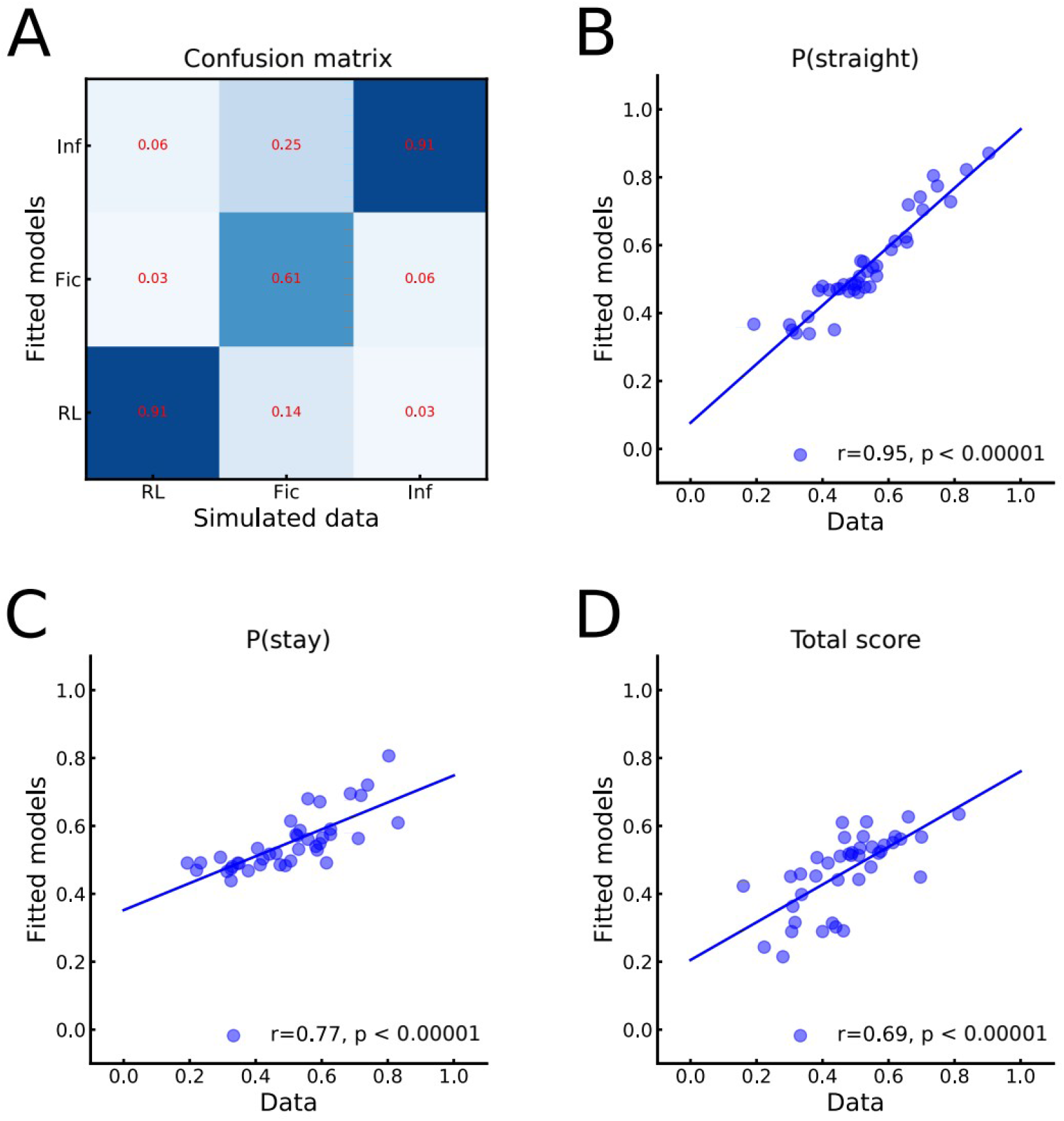
Evaluation of hyperparameter optimization for the value-based decisionmaking models. **A)** Confusion matrix summarizing the results of model recovery. All three models were fitted to 100 simulated subjects. Each cell represents the proportion of the simulated data that was best fitted by a certain model. Results showed moderate to strong recovery rates for all models. **B, C and D)** Each participant’s best fitting model was simulated again 10 times and the average probability of going straight, probability of repeating the previous action, and total scores was compared to actual participants’ data in order to validate the models. **B)** Correlation between the probability of going straight in participants’ data and in simulated data from each participant’s best fitting model. **C)** Correlation between the probability of repeating the previous action in participants’ data and in simulated data from each participant’s best fitting model. **D)** Correlation between total scores in participants’ data and in simulated data from each participant’s best fitting model.

**Supplementary Figure 3:**
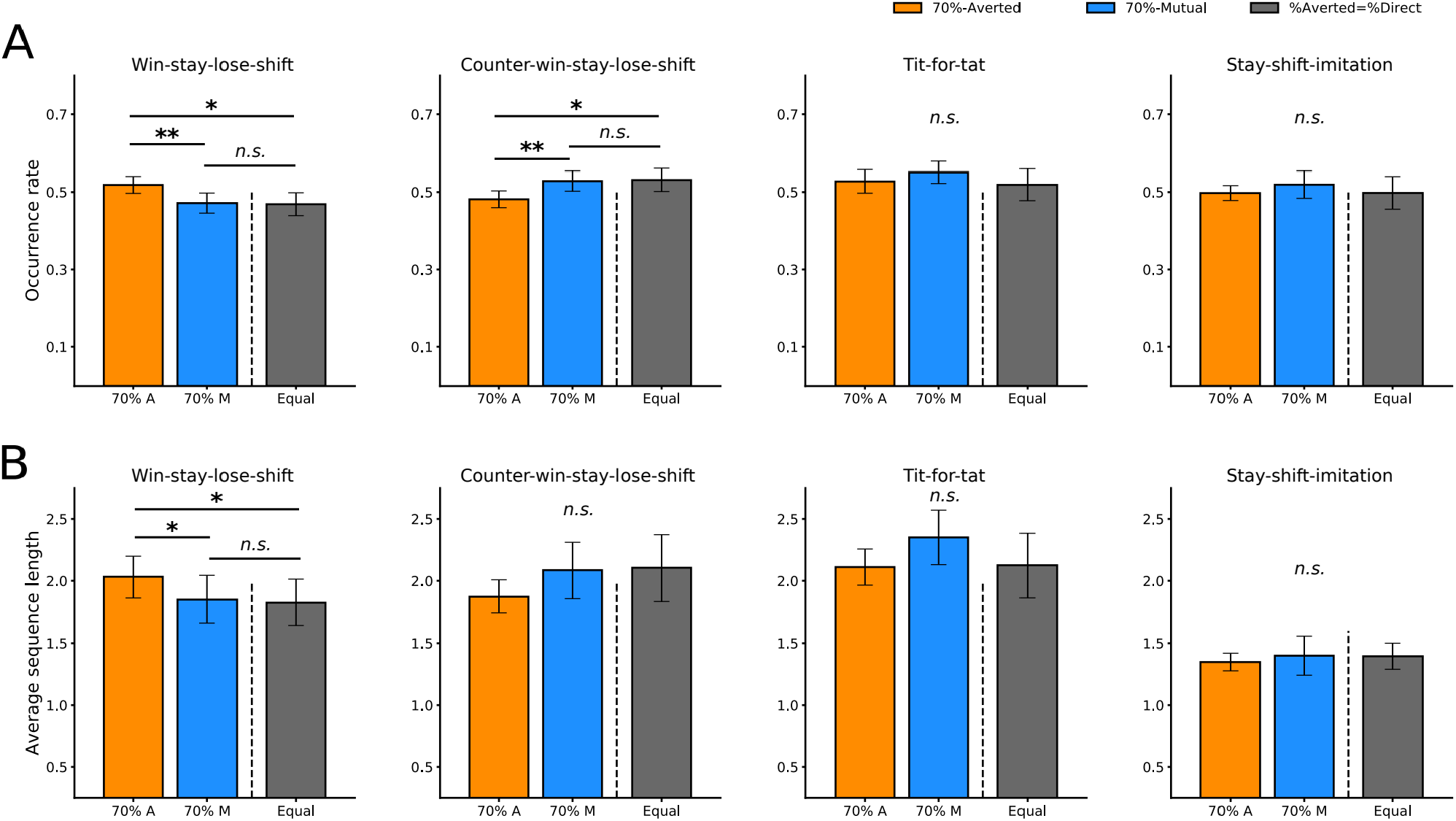
Patterns of self-oriented and other-oriented strategic behaviors in participants’ choice sequences. **A)** Occurrence rates of one self-oriented (win-stay-lose-shift, WSLS) and three other-oriented (counter-win-stay-lose-shift, C-WSLS; tit-for-tat, T4T; stay-shift-imitation, SS-Imit) patterns. 70% Averted and 70% Mutual conditions are taken from the main experiment’s data whereas the Equal condition is based on data from the pilot study. Occurrence rates of WSLS and C-WSLS in the 70% Mutual condition are similar to the Equal condition. They are significantly lower than 70 Averted in the case of WLSL and significantly greater in the case of C-WSLSL (70%A vs 70%M, U=108.5, p=0.006, 70%A vs Equal, U=100.0, p=0.017, Mann-Whitney test). No significant difference was found for T4T and SS-Imit. **B)** The average length of sequences of the WSLS pattern was significantly greater in the 70% Averted condition (70%A vs 70%M, U=120.0, p=0.015, 70%A vs Equal, U=114.0, p=0.045, Mann-Whitney test). No significant difference was found for C-WSLS, T4T and SS-Imit. N=20 for 70%A and 70%M and N=18 for Equal in A. Error bars represent 95% confidence intervals. * p < 0.05, ** p < 0.01. n.s., not significant at p > 0.05.

**Supplementary Table 1:** iCub’s verbal utterances in English (pilot study) and Italian (main experiment) which were randomly sampled 40% of the trials depending on the trial outcome in order to maintain participants’ engagement in the task

| Trial type | Utterance in English | Utterance in Italian |
| --- | --- | --- |
| Participant goes straight<br>iCub goes straight | “Oh no!” | “Oh no!” |
|  | “Ouch!” | “Ouch!” |
|  | “Too bad!” | “Peccato!” |
| Participant goes straight<br>iCub deviates | “Ohh” | “Ohh” |
|  | “Nice move” | “Bella mossa” |
|  | “Well done” | “Ben fatto” |
| Participant deviates<br>iCub goes straight | “Yess!” | “Sii!” |
|  | “Nice!” | “Evvai” |
|  | “You’re safe” | “Sei salvo” |
| Participant deviates<br>iCub deviates | “Good” | “Bene” |
|  | “We’re safe” | “Siamo salvi” |
|  | “Mmm” | “Mmm” |

**Supplementary Table 2:**
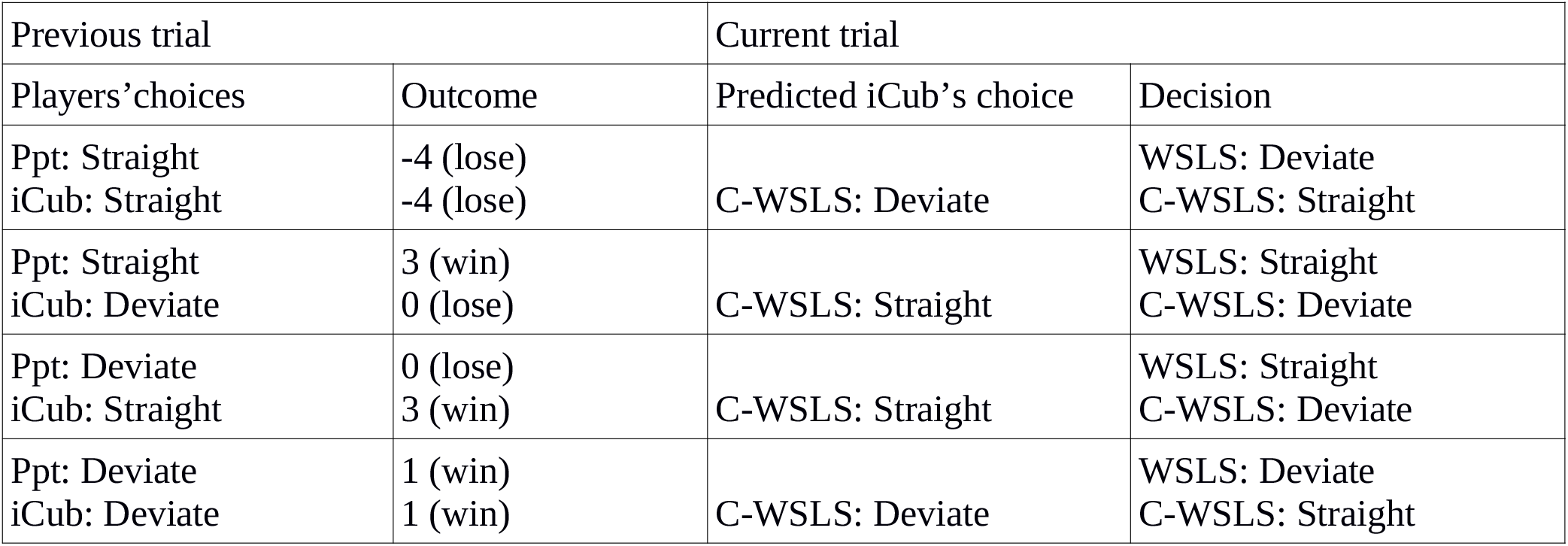
Detailed description of the player’s decisions as predicted by the win-stay-lose-shift (WSLS) and counter-win-stay-lose-shift (C-WSLS) strategies showing that these strategies lead to opposite choices in the context of our experiment.

**Supplementary Table 3:**
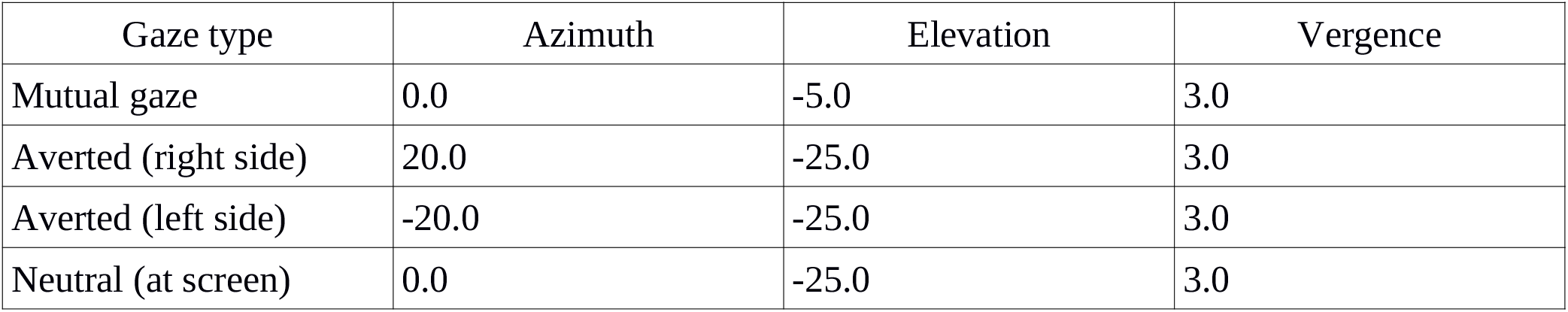
Azimuth, Elevation and Vergence (in degrees) in the absolute frame of reference fed to the robot gaze controller to direct eye movements to the desired positions.

| Gaze type | Azimuth | Elevation | Vergence |
| --- | --- | --- | --- |
| Mutual gaze | 0.0 | -5.0 | 3.0 |
| Averted (right side) | 20.0 | -25.0 | 3.0 |
| Averted (left side) | -20.0 | -25.0 | 3.0 |
| Neutral (at screen) | 0.0 | -25.0 | 3.0 |

## Notes

### Competing Interest Statement

The authors have declared no competing interest.

